# Human Sperm-Induced Cell-Cell Fusion Requiring JUNO (hSPICER): A paradigm shift to test sperm fertilizing potential

**DOI:** 10.64898/2026.05.07.723220

**Authors:** Nicolas G. Brukman, Maharan Kabha, Rivka Levi, Rinad Abbas, Shira Baram, Ronit Beck-Fruchter, Benjamin Podbilewicz

**Author notes:** Deceased on 23 June 2026.

## Abstract

Current evaluation of male fertility is largely based on indirect sperm parameters such as viability, concentration, morphology, and motility; however, each of these parameters, alone or combined, has been shown to have limited predictive value for higher fertilization rates. To address this problem, we introduce hSPICER (human SPerm-Induced CEll-cell fusion Requiring JUNO), an assay that evaluates sperm function based on their ability to induce fusion of somatic cells expressing human JUNO (hJUNO), the egg-specific sperm receptor. Similarly to our previous discovery in mice, we found that human sperm can fuse with somatic cells expressing hJUNO on their surface (pseudo-eggs) and promote content mixing between cells in culture, as measured using a split GFP system. The assay is sensitive, specific, and species-dependent, requiring hJUNO for optimal signal. We generated stable cell lines expressing hJUNO, enhancing reproducibility and sensitivity. We also show that hSPICER is compatible with cryopreserved sperm and consistent over different days. Importantly, hSPICER values correlate with in vitro fertilization outcomes of patients during fertility treatments, indicating its potential as a functional diagnostic tool. Beyond diagnostic uses, hSPICER establishes a platform to explore sperm fusion mechanisms and to screen for therapeutic compounds and interventions to treat low fertility, enhance fertilization, and develop non-hormonal contraceptives for males and females, as well as quality assessment of semen samples in fertility clinics and sperm banks.

## Introduction

Egg and sperm merge into a single cell, the zygote, in the culmination of a multi-step complex process known as fertilization (Yanagimachi, 1994). During the last steps of gamete interaction in mammals, the sperm plasma membrane protein IZUMO1 (Inoue et al., 2005) binds to its receptor on the oocyte membrane, named JUNO (Bianchi et al., 2014). This interaction is followed by gamete fusion, a process in which IZUMO1 functions as a unilateral fusogen, being required only on the sperm membrane (Brukman et al., 2023). Diverse gamete alterations before and during fertilization can lead to fertility defects.

Infertility affects 17.5% of the human population (World Health Organization, 2023), and a male factor is estimated to be a primary or contributing cause in approximately 50% of the cases (Agarwal et al., 2021). During the fertility evaluation of a male patient, a basic semen examination is performed, where different sperm parameters are examined: concentration, morphology, motility, and viability (World Health Organization, HRP, 2021). These results, while relatively easy to score, are often misleading as 12% of infertile male patients and only 60% of fertile men present normal sperm parameters (Boeri et al., 2021), evidencing the weak predictive value of these indirect variables. In light of the limited predictive value for fertilization outcomes of the conventional semen analysis, numerous specialized assays have been developed to evaluate specific functional aspects of sperm related to their capacity to fertilize an oocyte, including sperm-Zona Pellucida (ZP) binding (Oehninger et al., 2014) and penetration (de la Fuente et al., 2025) tests, the hyaluronan-binding assay (Pregl Breznik et al., 2013), DNA fragmentation (Fernández et al., 2003, 2005, 2011), the Hamster Oocyte Penetration (HOP) test (Yanagimachi et al., 1976; Aitken & Elton, 1986), and sperm membrane potential assessment (Molina et al., 2019), among others. Despite a myriad of possible assays, none of these methods is used routinely in clinics. Neither the American Urological Association nor the American Society for Reproductive Medicine currently recommends the regular clinical use of these sperm function tests, largely due to insufficient standardization, variability in threshold definitions, and a lack of robust, high-quality clinical evidence supporting their widespread applicability (Agarwal et al., 2021; Sanyal et al., 2024).

We have recently developed a new methodology to test sperm fertilizing function in mice based on their ability to induce the fusion of somatic cells expressing ectopically the egg-specific sperm receptor, JUNO (Brukman et al., 2024). When one sperm fuses with two adjacent cells, it bridges them and forms multinucleated syncytia in a process that is independent of *de novo* translation. We named this process SPICER (SPerm-Induced CEll-cell fusion Requiring JUNO), which mimics the final stages of natural fertilization and correlates with the ability of sperm to fuse with actual mouse oocytes (Brukman et al., 2024). We found that mouse sperm with higher fertilizing ability *in vitro* can induce the fusion of somatic cells more efficiently, judged by the increased levels of multinucleation and cytoplasmic mixing. The latter can be detected by the mixing of fluorescent markers (Avinoam et al., 2011; Brukman et al., 2023; Moi et al., 2022; Perez-Vargas et al., 2014; Podbilewicz et al., 2006; Valansi et al., 2017) or by the complementation of a split GFP system (Nakane & Matsuda, 2015).

Here, we found that SPICER can be applied to assess the fertilizing potential of human sperm (human SPICER, hSPICER). This novel methodology could complement current semen analysis protocols by providing a functional readout of an important component of sperm fertilizing potential. In general, hSPICER requires the transient transfection of a plasmid encoding for human JUNO (hJUNO), which adds more work and more variability to the assay. To overcome these caveats, we established modified cell lines that stably express hJUNO, enabling a robust, sensitive, and standardized methodology. Thus, hSPICER represents a powerful opportunity to evaluate sperm function with important applications in basic research, drug development, and medical treatments of infertility.

## Results

### Human sperm can fuse to somatic cells expressing hJUNO

We have recently shown that mouse sperm can fuse to cells expressing the IZUMO1 receptor (JUNO) localized to the plasma membrane. To evaluate if human sperm can fuse to somatic cells expressing hJUNO we transfected Baby Hamster Kidney (BHK) cells in culture with a plasmid encoding for hJUNO or a control plasmid (negative control) together with a reporter protein linked to GFP that binds DNA (GFP-MBD) (Yamagata et al., 2005). BHK cells facilitate visualization of the cytoplasm due to their strong adherence and spread morphology. We obtained semen samples from normozoospermic donors, isolated the sperm using density gradient centrifugation, capacitated, and incubated them with these JUNO-BHK and control BHK cells for 6 h. When hJUNO was present, we detected fusion events between the somatic cells and the sperm judged by the transference of the GFP-MBD to the sperm nucleus (Figure 1). This result indicates that human sperm can fuse with somatic cells ectopically expressing hJUNO which can mimic human oocytes (pseudo-eggs).

**Figure 1:**
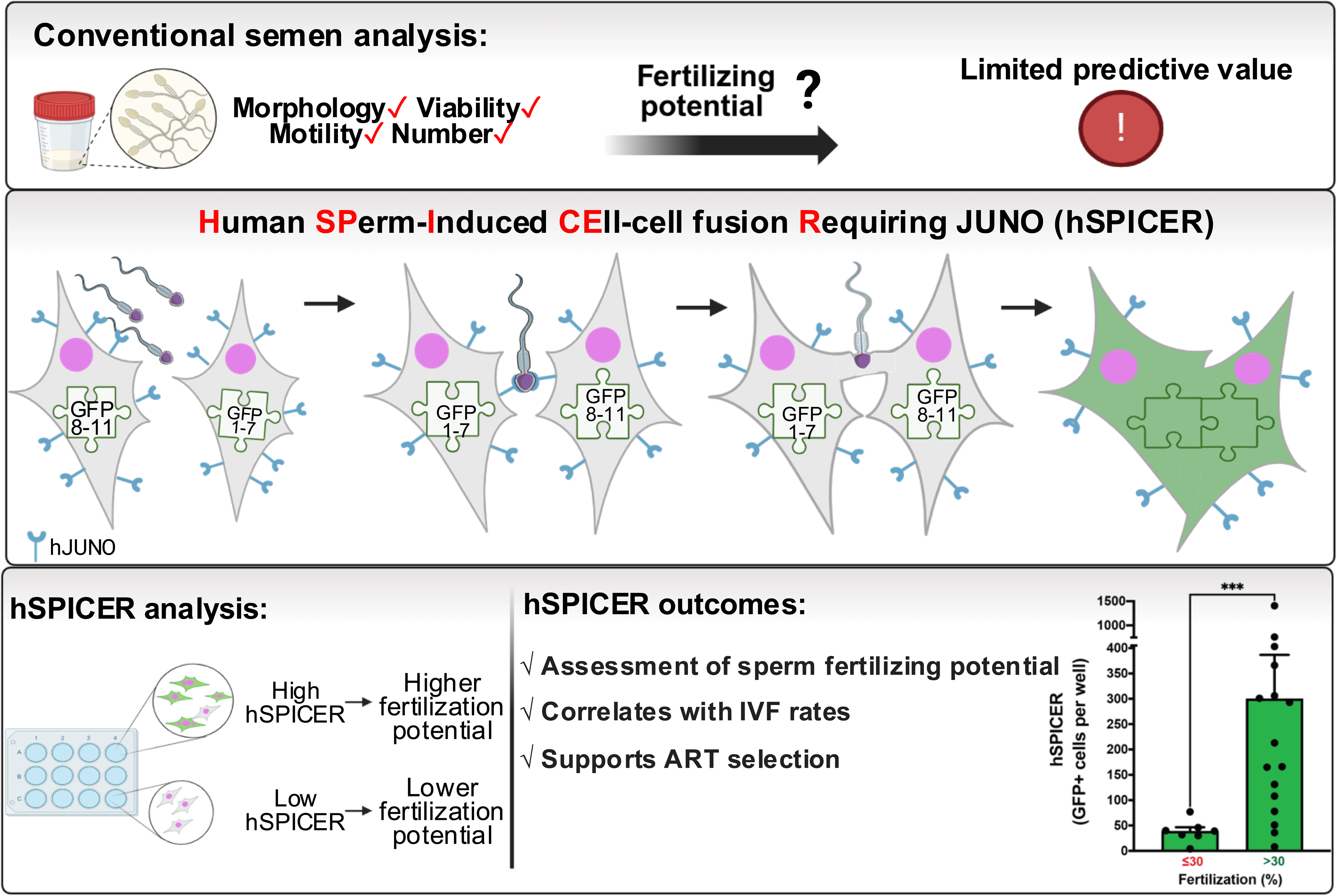
Human sperm can fuse to somatic cells. (A) BHK cells were transfected with pCI::H2B-RFP or pCI::hJUNO::H2B-RFP vectors together with the plasmid pcDNA3.1-EGFP-MBD-nls. The cells were later co-incubated with human sperm for 6 h, fixed, and stained with DAPI and Mitotracker. (B) Fusion was evaluated as the presence of GFP-MBD signal (green) inside the sperm heads (arrowhead). Bound sperm, without GFP-MBD staining (unfused), are pointed with arrows. The contour of a tri-nucleated cell is marked with a dashed line. Scale bar, 20 µm. An amplified image of the squared region is presented in the bottom row.

### Human sperm induce cell-cell fusion of somatic cells expressing hJUNO

SPICER is a test developed to study the fertilizing ability of sperm using somatic cells that fuse if they express surface JUNO because the sperm can simultaneously merge with two or more somatic cells (Brukman et al., 2024). SPICER was originally developed using sperm and JUNO from mice. To study whether somatic cell-cell fusion occurs if human sperm is used (hSPICER), we employed two cell lines of Human Embryonic Kidney (HEK) cells engineered to express a different half of a green fluorescent protein (split GFP) (Brukman et al., 2024) that were mixed and transfected with a plasmid encoding for hJUNO or a control plasmid (negative control). HEK cells were selected for their ease of culture, high transfectability, and our previous successful use of this cell line in the mouse SPICER system (Brukman et al., 2024). Only following the fusion of the cells of the two populations, the assembly of the two halves completes the GFP, and the fluorescent syncytia can be quantified (Figure 2A). If human sperm can induce the fusion of HEK cells, then we expect that GFP will be reconstituted, and we will detect cells with green fluorescence. Sperm cells obtained from normozoospermic donors were capacitated and then incubated with the HEK cells for 24 h. We detected GFP-positive cells when the cells were transfected with a plasmid encoding for hJUNO and incubated with sperm from different normozoospermic donors, but not when a control plasmid was used or when the sperm were excluded (Figure 2B, C; Table S2). After accounting for the transfection efficiency (47.5 ± 5.5 %, n=3) and the number of cells in the plate at the moment of co-incubation (8.7 ± 0.9 x10^5^ cells, n=3), both determined independently under the same experimental conditions, an average of approximately 600 GFP-positive cells per well corresponds to an estimated 0.15 % of the hJUNO-expressing cell population. Given the relatively low frequency of fusion events in the human assay compared with the mouse assay, results were expressed as the absolute number of GFP-positive cells per well to provide a more sensitive comparative readout. We noted a range of two orders of magnitude between different donors in the number of GFP-positive cells, suggesting that hSPICER can detect variability among the male population (Figure 2C). The results suggest that human sperm can induce the fusion of the somatic cells and show that hSPICER could represent a novel research tool to study the sperm potential for fertilization in the presence of pseudo-eggs (somatic cells expressing hJUNO).

**Figure 2:**
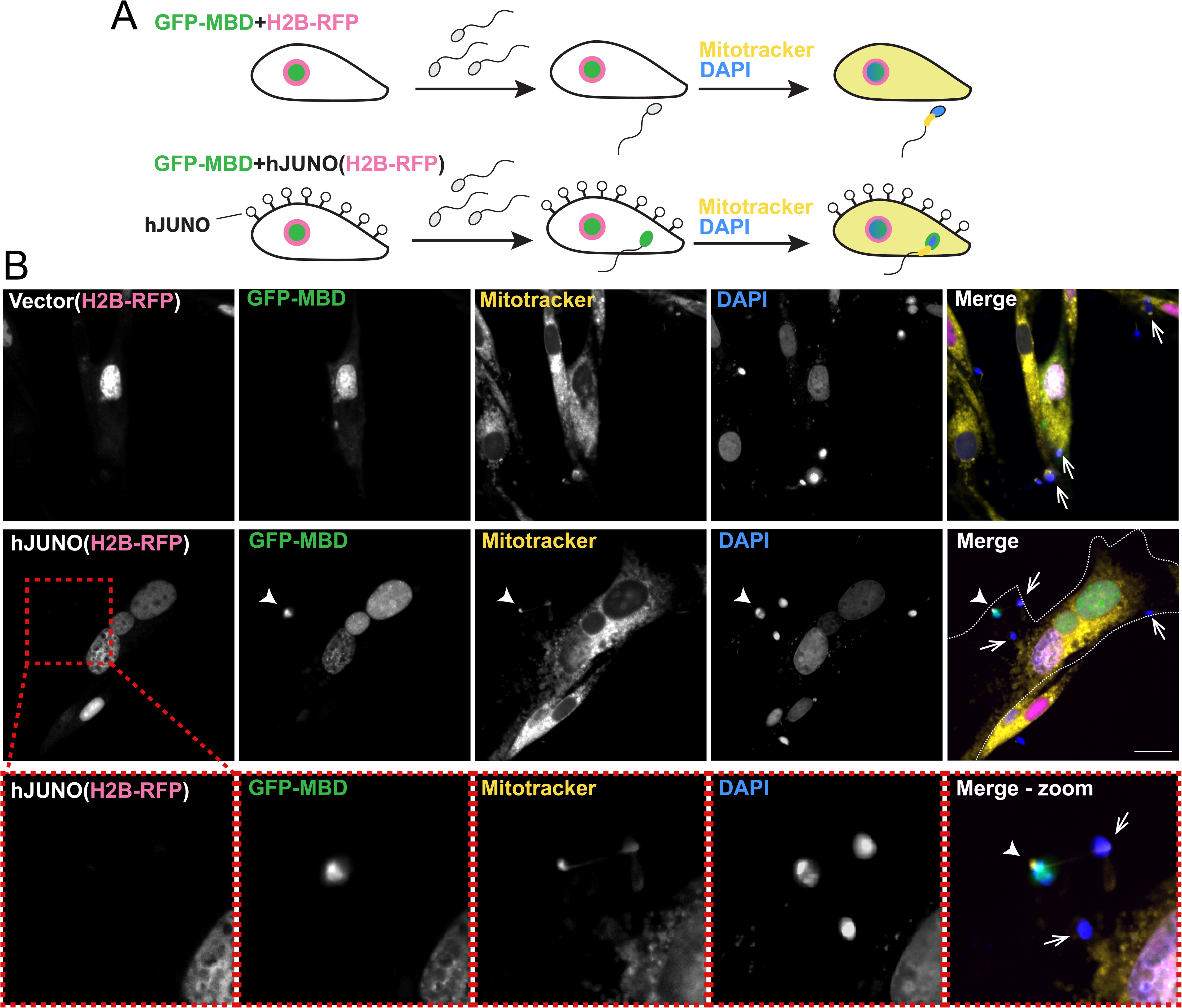
Human sperm mediate fusion of human cells expressing hJUNO. (A) HEK293T cells stably expressing a split GFP (GFP1-7 or GFP8-11, corresponding to DSP1-7 and DSP8-11) were mixed and transfected with pCI::H2B-RFP or pCI::hJUNO::H2B-RFP plasmids. Where indicated, the cells were later co-incubated with capacitated human sperm for 24 h, and fixed. (B) Representative images for each treatment showing GFP syncytia (assembled GFP in fused cells) and H2B-RFP (magenta nuclei). Arrows indicate two representative GFP-positive cells observed in the hJUNO + sperm condition. Scale bar, 50 µm. (C) Quantification of content mixing experiments. The extent of fusion was determined by counting the number of GFP-positive cells per well. Bar chart showing individual experiment values and means ± SEM of 7 independent experiments using sperm from different donors (Table S2). Comparisons by one-way ANOVA followed by Dunnett’s test. ns = non-significant, *p<0.05.

To study the kinetics of hSPICER we used GFP reconstitution to approximate cell-cell fusion levels over a 24 h period. We found that, following a lag of about 2 h, there is an apparent linear increase in cell-cell fusion induced by sperm, followed by a stationary phase, with the maximum signal reached after 8 h of co-incubation (Figure 3A).

**Figure 3:**
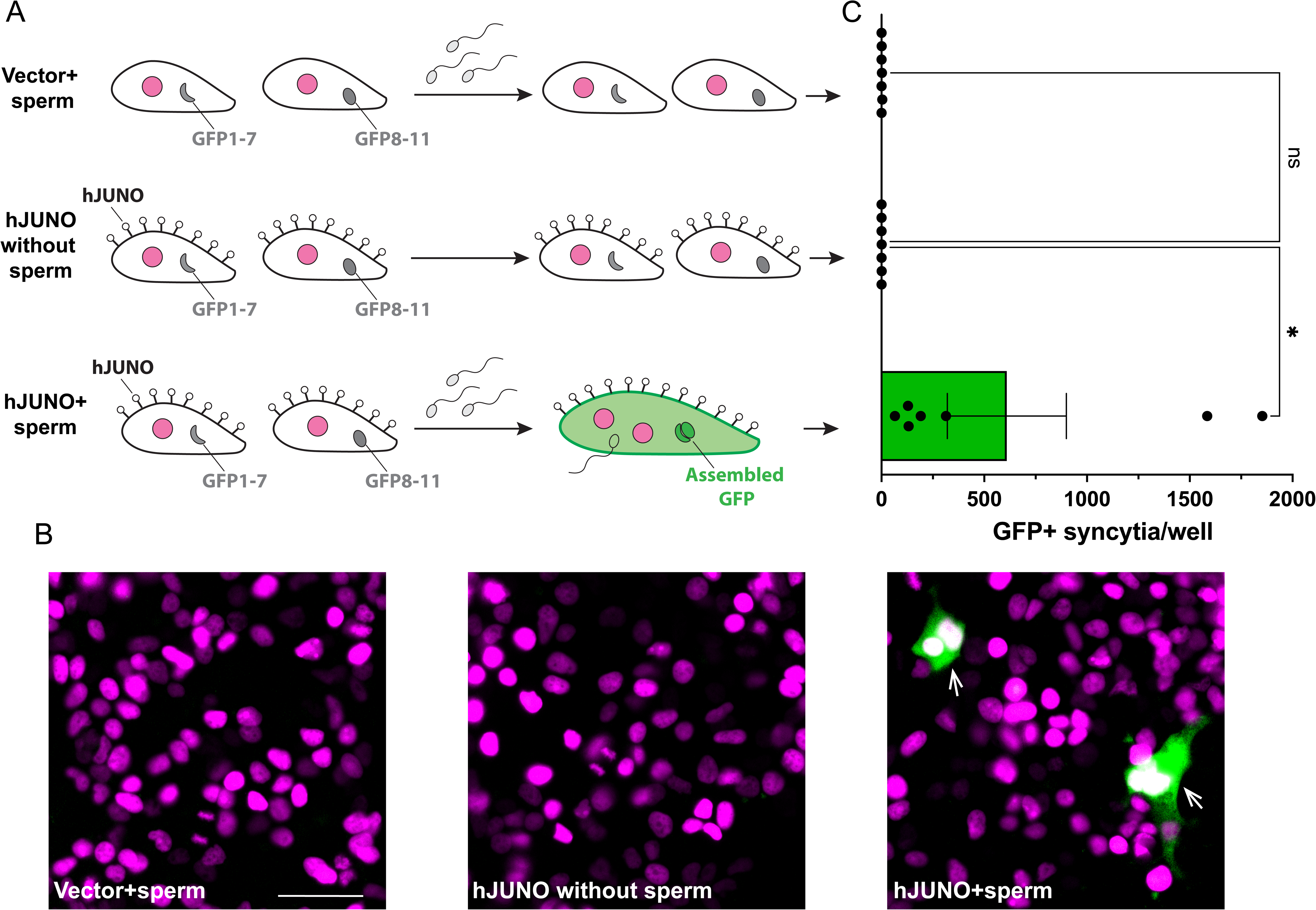
Efficient hSPICER requires optimal time, sperm concentration, and human JUNO. (A-B) HEK293T cells stably expressing a split GFP (DSP1-7 or DSP8-11) were mixed and transfected with pCI::H2B-RFP or pCI::hJUNO::H2B-RFP vectors. Later, the cells were co-incubated for different times with 10^7^ human sperm for various amounts of time (A) or with varying amounts of sperm for 24 h (B). The extent of fusion was determined by counting the number of GFP-positive cells per well relative to the maximum signal (24 h (A) or 10^7^ sperm (B), and therefore have no error bars). Graphs show means ± SEM of 5 independent experiments. (C) HEK293T cells were transfected with plasmids coding for mouse, hamster or human JUNO or the empty vector, co-incubated with 10^7^ human sperm for 24 h and the number of GFP-positive cells quantified. The values are per well relative to the signal of human JUNO. Comparisons by one-way ANOVA followed by Dunnett’s test. ns = non-significant, ***p<0.001, ****p<0.0001. The 95% confidence intervals for the normalized values at 8 h in (A) and with 5 × 10 sperm in (B) included 100%, indicating that no statistically significant difference from the respective reference conditions was detected.

To determine the optimal amount of sperm required for hSPICER we used increasing amounts of sperm and estimated the levels of sperm-induced cell-cell fusion following a 24 h incubation. We found that the optimal amount of sperm for the assay is between 0.5-1×10^7^ sperm cells per well (Figure 3B). Last, to study the species-specificity of this assay, we used pseudo-egg somatic cells expressing mouse and hamster JUNO and compared them with hJUNO. We found that hSPICER requires a species-matching JUNO, as mouse or hamster JUNO gave only around 2% of the GFP signal achieved using the human protein (Figure 3C). Thus, we have optimized the hSPICER assay and showed that 8 h is the minimal time to obtain maximum signal when using 0.5-1×10^7^ sperm per well.

### Stable cell lines expressing hJUNO support a robust and reproducible hSPICER

To standardize and simplify the assay, we next aimed to generate cell lines stably expressing hJUNO. This eliminates the requirement for a transfection step, and provides a stronger, uniform expression. To facilitate such optimization, we first investigated the possibility of adding tags to the ectodomain. We explored three options (Figure 4A): adding a flag tag between the signal peptide and the ectodomain (N-ter flag); a flag tag between the ectodomain and the Glycosylphosphatidylinositol (GPI) signal (C-ter flag); and replacing the GPI signal by the transmembrane domain of IZUMO1 together with a V5 tag (TMD-V5). This last construct was designed based on our previous observation that the TMDs of the fusogens Fsx1 from archaea and Arabidopsis HAP2 can be replaced by GPI anchors while retaining fusogenic activity (Moi et al., 2022). A tag introduced downstream of the TMD, allows detection of the protein without introducing a tag into its extracellular domain, where it could potentially interfere with its function. We then determined the expression of the different variants in HEK cells by immunostaining in permeabilized and non-permeabilized cells. We found that while both flag-tagged variants were expressed at similar levels when a permeabilization step was performed, in non-permeabilized cells only the C-ter flag was strongly detected on the plasma membrane, suggesting that this variant can be trafficked to the surface more efficiently (Figure 4B). The V5 tag of the TMD-V5 confirmed the protein is highly expressed in permeabilized cells, however we were unable to detect the transmembrane protein on the surface in non-permeabilized cells due to its intracellular placement (Figure 4B). To establish whether the tagged hJUNO variants produce pseudo-eggs compatible with hSPICER, we compared their performance in HEK cells against wildtype hJUNO. We found that the C-ter flag is as functional as the untagged hJUNO (Figure 4C) and was the best candidate for generating stable lines.

**Figure 4:**
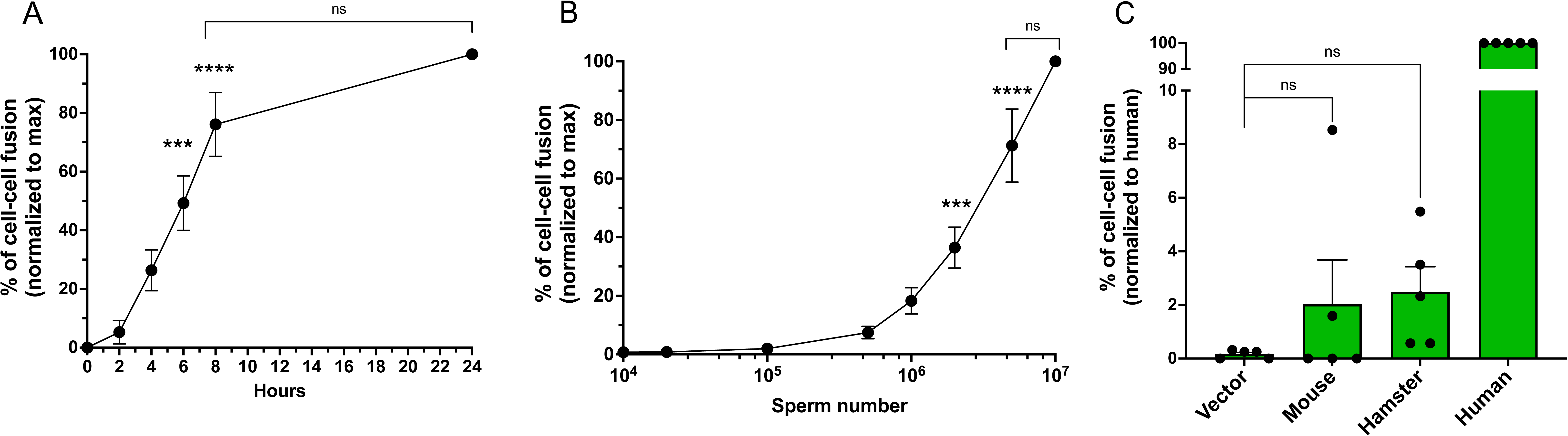
A C-terminally tagged hJUNO is functional. (A) Representation of wildtype hJUNO including the signal peptide (SP), the ectodomain, and the GPI signal (GPI). Two different variations of JUNO containing a flag tag in the N– or C-terminal part of the ectodomain are represented. In addition, the GPI was replaced by IZUMO1’s transmembrane domain (TMD) followed by a V5 tag. (B) Representative micrographs showing immunostaining (in white) for permeabilized and non-permeabilized HEK cells, using as primary antibody anti-flag (for N-ter and C-ter variants) or anti-V5 (for TMD-V5). H2B-RFP (magenta nuclei) are included. Scale bar, 20 µm. Immunofluorescence analysis showed that the N-flag, C-flag, and TMD-V5, hJUNO constructs were expressed in 67.5%, 97.8%, and 96.9%, of the transfected permeabilized cells, respectively, while cell-surface staining in non-permeabilized cells was detected in 25.0%, 80.5%, and 3.25%, of transfected cells. (C) HEK293T cells stably expressing a split GFP (GFP1-7 or GFP8-11) were mixed and transfected with pCI::H2B-RFP (Vector) or pCI::hJUNO::H2B-RFP vectors (wildtype (WT), C-terminal flag (C-ter; hJUNO-flag), N-terminal flag (N-ter; flag-hJUNO), or transmembrane anchored (TMD-V5; hJUNO-TMD-V5)). The cells were later co-incubated with human sperm for 8 h, and fixed. The extent of fusion was determined by counting the number of GFP-positive cells per well and normalized to the wildtype. Bar chart showing individual experiment values and means ± SEM of 3 independent experiments. ns = non-significant, *p<0.05.

To generate stable lines expressing hJUNO, the sequence of *hJUNO-flag* (in the C-terminal part of the ectodomain), was cloned into a lentiviral vector that co-expresses mCherry (Figure 5A). Lentiviral particles containing the sequence for *hJUNO-flag* were transduced into HEK cells stably expressing the split GFP constructs. The success of the transduction was confirmed by evaluating the fluorescence of mCherry and the expression of hJUNO-flag by immunostaining (Figure 5B). The final expression levels were quantified and compared to HEK cells transiently transfected with the plasmid encoding for hJUNO-flag and H2B-RFP (Figure 5B, C). We found that almost all the cells in the stable lines expressed the sperm receptor (Figure 5C). Next, the ability of these stable lines to facilitate hSPICER was measured relative to transient hJUNO transfection. 2×10^6^ sperm cells per well was selected as the lowest number required to obtain a significant signal (Figure 3B), thereby minimizing the amount of material needed. We found a 2.6-fold increase in the efficiency of hSPICER when the new stable lines were incubated with human sperm compared to transient expression (Figure 5D). The expression of the protein in the stable cell lines was analyzed by Western blotting, obtaining a band of approximately 30 kDa, corresponding to the glycosylated, flag-tagged JUNO (Figure 5E). The generation of these cell lines is an important step in the development of a sensitive, simple, and reproducible male infertility diagnostic tool.

**Figure 5:**
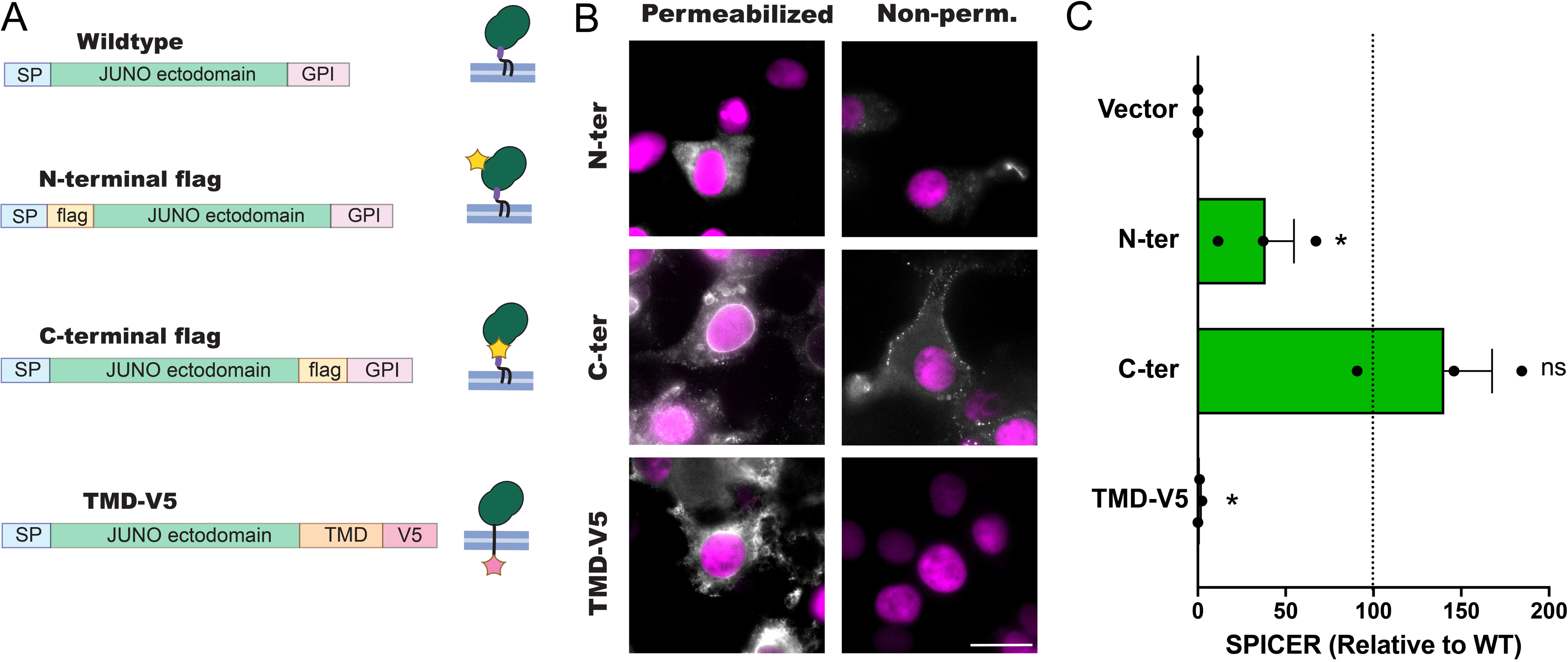
Cell lines stably expressing hJUNO-flag can mediate hSPICER more efficiently. (A) HEK293T cells were transfected with the empty pCSCW::IRES::mCherry plasmid or bearing the sequence for hJUNO with a C-terminal flag tag. Immunofluorescence was performed using anti-flag antibodies. DAPI-stained nuclei (blue) and mCherry (magenta) are shown. Scale bar, 20 µm. (B) HEK293T cells stably expressing a split GFP (DSP1-7 or DSP8-11) were transduced with the lentiviral vector pCSCW::hJUNO-flag::mCherry. The levels of expression were tested by immunostaining of permeabilized cells using anti-flag antibodies and compared to those of HEK293T cells transiently transfected with pCI::hJUNO-flag::H2B-RFP. The immunostaining signal of flag (green), H2B-RFP or mCherry signals (magenta), nuclei staining (blue), and merge are shown. Scale bar, 50 µm. (C) Quantification of expression levels as the percentage of hJUNO expressing cells (with signal for Flag) over the total cells. At least 200 cells in 4 independent experiments were counted. The bar chart shows individual experiment values and means ± SEM. Comparisons by one-way ANOVA followed by Dunnett’s test. ****p<0.0001. (D) HEK293T cells stably expressing a split GFP (DSP1-7 or DSP8-11) transiently or stably expressing hJUNO-flag were mixed. The cells were later co-incubated with 2×10^6^ human sperm. The extent of fusion was determined by counting the number of GFP-positive cells per well and normalizing them to the transient transfection. A control without sperm is included. Bar chart showing individual experiment values and means ± SEM of 10 independent experiments. The mean of the “Stable” was analyzed by one-sample t test vs. 100. ***p<0.001. (E) Total protein extracts from wild-type HEK293T (WT) or DSP1-7 and 8-11 expressing hJUNO were evaluated for the presence of hJUNO-flag (arrowhead) by western blotting using an anti-flag antibody. Tubulin was used as loading control (bottom panel). The position of the molecular weight standards is indicated in kDa on the left. Representative blots are shown.

Following the establishment of stable cell lines expressing hJUNO-flag and split-GFP to facilitate robust hSPICER, we proceeded to evaluate the system as a potential predictor of sperm function. For this, we shifted from the sequential capacitation and co-incubation used for the initial characterization of the system to concomitant capacitation and co-incubation with the cells, thereby more closely mimicking clinical *in vitro* fertilization (IVF) protocols. First, we evaluated whether the assay could be performed using cryopreserved sperm, thereby providing greater flexibility in experimental timing. For this, sperm samples were tested fresh or after a freezing/thawing cycle. As expected, the results show a decrease in the fusion potential of thawed sperm, however, this reduction is less than 40% (Figure 6A), and there is a significant correlation with the values in the fresh samples (Figure 6B). We also evaluated the reproducibility of the assay by testing frozen aliquots of the same sperm samples in different weeks. We observed that the values obtained in different experiments for frozen aliquots of the same samples were similar (Figure 6C). Finally, to determine the functional significance of hSPICER we evaluated sperm samples from patients undergoing conventional IVF treatment and compared the levels of fusion obtained from sperm with low fertilization rates (<30% of fertilization) to those with higher fertilization rates (Figure 6D-F). This threshold is routinely used by the IVF Unit at HaEmek Hospital, based on previously established definitions of fertilization failure (Montag et al., 2012; Murugesu et al., 2017; Ruan et al., 2023). While the Total Motile Cells (TMC) parameter, used in standard sperm analysis, was similar between groups (Figure 6D), the results show that hSPICER values are significantly higher for the sperm samples that resulted in higher fertilization rates in IVF (Figure 6E, Table S3), suggesting that hSPICER can be a good predictor of conventional IVF outcomes. Receiver Operating Characteristic (ROC) curve analysis showed that hSPICER discriminated between samples associated with <30% and ≥30% IVF fertilization, with an area under the ROC curve (AUC) of 0.902 (95% CI, 0.773-1.000; p=0.0026). Using a cutoff of 77.5 GFP+ cells per well, the assay achieved 100% sensitivity and 81.3% specificity. In addition, hSPICER did not correlate with TMC (Figure 6F), suggesting that these two parameters are independent functional indicators. In summary, we have tested hSPICER using stable lines expressing hJUNO and sperm from different donors. We found high reproducibility and accuracy for the results using different donors and a large variability in the fertilization potential between individuals. Moreover, we uncovered that cryopreserved human sperm retains hSPICER functionality, although levels of fusion are reduced by 40%. The good correlation between results obtained in the IVF clinic and hSPICER levels suggests that this method efficiently reveals the fertilizing ability of human sperm.

**Figure 6:**
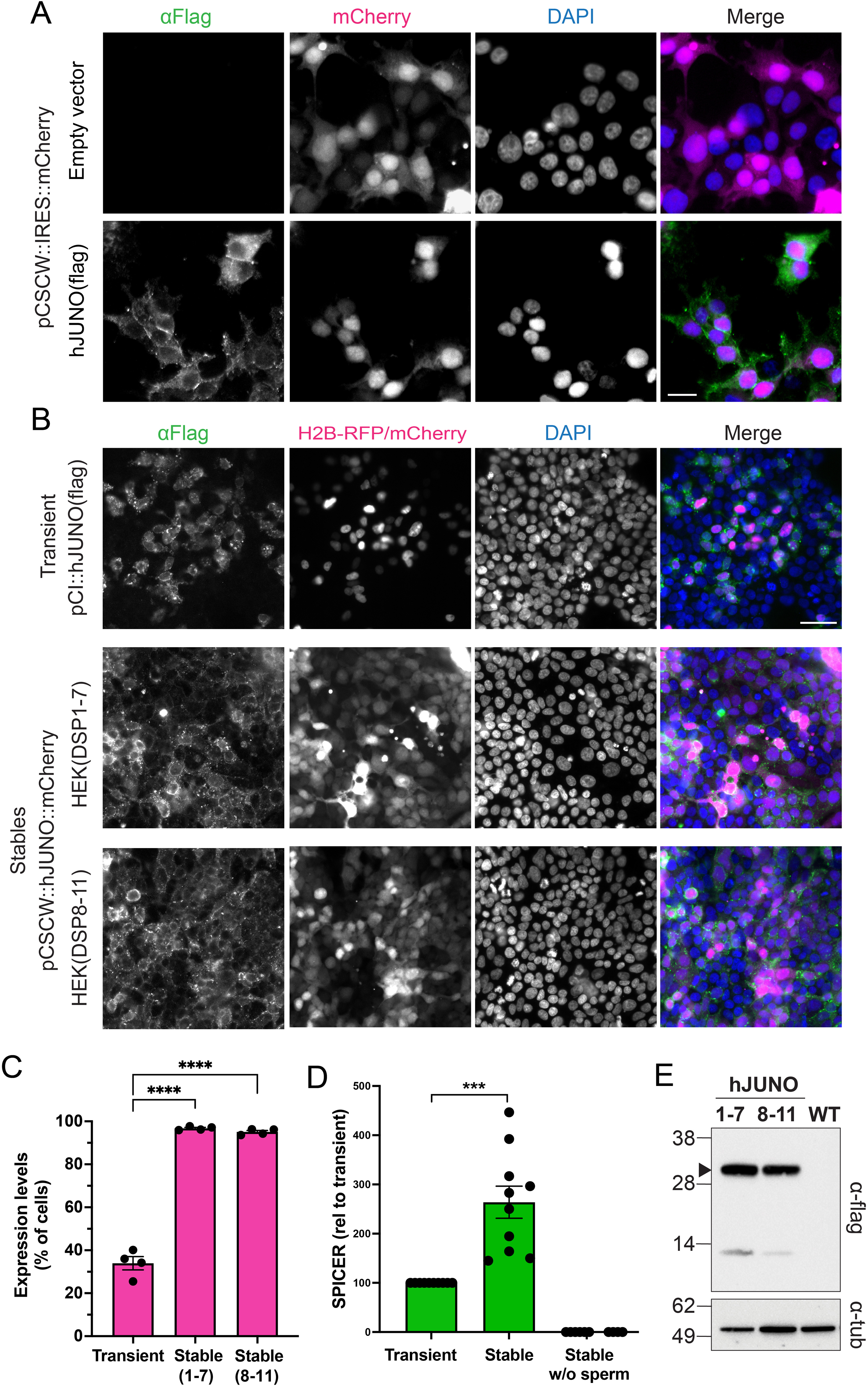
hSPICER is reliable, reproducible, can be used with frozen samples and correlates with the fertilizing activity of sperm during IVF treatments. (A) Quantification of hSPICER as the number of GFP-positive cells after a content mixing experiment using 2×10^6^ fresh (Fresh) or frozen/thawed (“Frozen”) sperm from the same semen sample. A control without sperm was included. The bar chart shows 7 individual experimental values and means relative to “Fresh” ± SEM. The mean of the “Frozen” sperm was analyzed by one-sample t-test vs. 100. **p<0.01. (B) The raw values of the experiment in (A) are plotted. A Pearson correlation analysis was performed, and the “r”, “R^2^” and p values are shown in the graph. (C) Quantification of hSPICER as the number of GFP-positive cells after content mixing experiments performed in different weeks, for frozen aliquots from the same semen sample. n=4. (D and E) Total Motile Cells in semen (TMC, D) and hSPICER (E) values of samples from patients who obtained low (less than or equal to 30%, n=7) or high (more than 30%, n=16) fertilization rates during fertility treatments. The bar chart shows individual experiment values and means ± SEM. Comparison made by Mann-Whitney test. ***p<0.001. (F) hSPICER as the function of TMC. Each dot corresponds to a different patient (see Table S3). A Pearson correlation analysis was made, and the “r”, “R^2^”, and p values are shown in the graph.

## Discussion

Our results indicate that human sperm obtained from normozoospermic donors can induce the fusion of somatic cells expressing human JUNO in a process called hSPICER. This process occurs in a reproducible and robust way, and can be readily measured, allowing various applications. In basic research, it enables the study of the fertilization machinery of the sperm without oocytes, which are normally a limiting material. In translational approaches, it can assist in the screening of compounds that can enhance egg-sperm fusion and could work as new fertility treatments, or that block fusion and could represent interesting male and female contraceptive alternatives. In medicine, hSPICER could evaluate the male contribution to a couple’s infertility and predict the suitability of different Assisted Reproductive Technologies (ARTs) by evaluating the fusogenic ability of sperm. In this context, conventional IVF requires fertilization-competent sperm capable of completing gamete interaction and fusion, whereas intracytoplasmic sperm injection (ICSI) bypasses these steps (Wang et al., 2024).

Sperm cells must undergo several processes to be able to fuse with the egg, including maturation, capacitation, and acrosome reaction (Gervasi & Visconti, 2016; Gierke et al., 2026; Hirohashi & Yanagimachi, 2018). Among the cellular changes that occur during these events, IZUMO1 relocalizes to the fusogenic region in the sperm head (Satouh et al., 2012). hSPICER provides a unique metric to the final facilitation of fusion, thereby encompassing most of previous steps which are a prerequisite for this process to occur. Successful hSPICER performance therefore indicates the efficiency of most previous cellular requirements leading up to the moment of fusion, including sperm capacitation, acrosome reaction, and binding. It should be noted, however, that the co-incubation time used in hSPICER is longer than the timeframe of gamete interaction observed in time-lapse studies of conventional IVF (Mio et al., 2012). This difference may reflect the time required for the accumulation of fusion events, cytoplasmic content mixing, and split-GFP complementation to generate a robust readout. As opposed to the traditional HOP (Yanagimachi et al., 1976; Aitken & Elton, 1986), our test does not require the use of experimental animals or the laborious collection of oocytes. However, hSPICER requires cell culture, which should be considered for its implementation in diagnostic laboratories. Here, we employed a lentiviral vector to establish cell lines with an integrated plasmid that encodes for hJUNO. In this way, we increased the percentage of cells expressing the transgene (Figure 5B, C), resulting in an improved sensitivity of the assay (Figure 5D) and reduced manipulation during the experiment. In addition, the higher consistency of levels of expression is translated into more reproducible results, and the fact that the transfection step is no longer required makes the methodology simpler, cheaper, and faster. These cells can be shared among different diagnostic laboratories, improving the standardization of the assay. Based on all this, this system could be applied in the future for screening of molecules that enhance or inhibit sperm function at any of the steps required for sperm fusion (as stated above, from capacitation to membrane merger). The use of a cell-based assay that relies on a simple readout sets the basis for an increased throughput; for instance, the counting of the GFP-positive cells can be automated using AI-based microscopic analyses, by flow cytometry or the fluorescence can be measured with a plate reader. Furthermore, the DSP system contains a second split protein, a split luciferase (Ishikawa et al., 2012; Nakane & Matsuda, 2015), enabling the additional detection of luminescence as hSPICER readout that could provide flexibility depending on the equipment available in different laboratories.

The results indicate that the heterologous expression of hJUNO is sufficient to enable the fusion of the sperm to somatic cells (Figure 1), consistent with previous observations in mice (Brukman et al., 2023, 2024). Considering that JUNO expression alone does not induce fusion (Figure 2; Brukman et al., 2023, 2024), it seems more likely that the sperm binding to this receptor is a prerequisite for membrane merger rather than being part of fusion itself. Furthermore, this does not exclude the possibility that other molecules from the egg side, which could also be expressed in these somatic cells, are required for gamete fusion. One example is the egg CD9, which was the first gamete molecule shown to be essential for egg-sperm interactions (Kaji et al., 2000; Le Naour et al., 2000; Miyado et al., 2000) and is widely expressed in different tissues, with its endogenous presence is also reported in HEK cells (Duke et al., 2025). In this sense, hSPICER represents a promising approach to studying these molecules in the absence of oocytes. From the sperm side, many proteins have been reported to be essential for gamete fusion: IZUMO1, SPACA6, TMEM81, TMEM95, DCST1/2, SOF1 and FIMP (Inoue et al., 2005; Barbaux et al., 2020; Noda et al., 2020; Lamas-Toranzo et al., 2020; Fujihara et al., 2020; Inoue et al., 2021; Noda et al., 2022; Elofsson et al., 2024; Deneke et al., 2024). The exact roles of each of them during the last steps of fertilization remain unclear. IZUMO1 has been extensively shown to interact with JUNO on the oocyte mediating gamete binding (Bianchi et al., 2014; Matsumura et al., 2022), however, based on its ability to mediate somatic cell-cell fusion in vitro, it is plausible that IZUMO1 is involved in the post-binding fusion as well (Brukman et al., 2023). This activity was comparable to other eukaryotic fusogens (e.g., EFF-1, AFF-1, and GCS1/HAP2) and lower than their viral counterparts (Podbilewicz et al., 2006; Avinoam et al., 2011; Perez-Vargas et al., 2014; Valansi et al., 2017; Kumar et al., 2022), which can explain why other studies failed to detect fusion of somatic cells expressing IZUMO1 and JUNO (Bianchi et al., 2014; Bianchi et al., 2024). Future work will determine whether IZUMO1 mediates gamete fusion in vivo and if the other essential sperm factors are required for efficient IZUMO1 activity. We anticipate that the human SPICER assay may provide a useful platform to investigate the function of their human orthologues during gamete fusion in the absence of oocytes.

Our improved methodology has been shown to be efficient with frozen sperm samples (Figure 6A, B) and with reproducible results over time. This is not only convenient to better control the time of analysis, but also indicates that hSPICER can be used as a quality control test for cryopreserved sperm from sperm banks and in fertility clinics. More importantly, as opposed to the classical TMC parameter, we were able to detect lower levels of hSPICER in patients who showed fertilization failure in conventional IVF treatment (Figure 6E, Table S3). This result supports that hSPICER is a reliable read-out of the sperm fertilizing potential. In this experiment, however, we noticed that two patients with higher fertilization rates induced levels of cell fusion comparable to those of the group with low fertilization rates (<50 GFP-positive cells per well). This could be explained by a lower cryotolerance of these particular samples (Holt, 2000), with these particular samples being more susceptible to cryopreservation, as the IVF was performed with the fresh sample and hSPICER after thawing. On the other hand, there is a patient with low fertilization rates but relatively high hSPICER levels, which could be explained by a female factor, such as mutations on JUNO-coding gene (Allingham & Floriano 2021), or sperm defects that are not fully reconstituted in hSPICER, such as interactions with the ZP, which has to be penetrated by the sperm in the conventional IVF. Indeed, defects in CatSper function can compromise sperm penetration of the ZP and result in fertilization failure despite otherwise preserved sperm parameters (Young et al., 2024). Interestingly, a recent report uncovered a late function for CatSper-mediated sperm motility in gamete fusion (González et al., 2026). While hSPICER may not identify sperm penetration defects in CatSper mutant sperm, it is expected that gamete interactions leading to oocyte-sperm fusion will be detected by our assay. Other possible defects that are not reconstituted in hSPICER are those associated with oocyte activation, such as alterations associated with the sperm-specific Phospholipase C Zeta1 (PLCZ1) (reviewed in Parrella et al., 2024). Despite these inherent limitations of the assay, a cutoff of 78 GFP+ cells per well showed 100% sensitivity and 81% specificity for discriminating samples according to IVF outcome. However, this threshold was derived from the present cohort and requires validation in larger, independent cohorts before it can be considered for clinical decision-making.

Sperm functional assays have been of interest to discriminate fertile from subfertile males and to predict the outcomes of ARTs (Oehninger et al., 2014). Unlike conventional semen analysis, several functional assays evaluate specific sperm interactions with components of the cumulus–oocyte complex, such as binding to hyaluronic acid (Huszar et al., 2003). More recently, recombinant JUNO-coated beads have been developed to assess the ability of cattle and human sperm to bind to the oocyte receptor (Cots-Rodríguez et al., 2026; Hamze et al., 2020). However, these assays evaluate sperm binding to an isolated receptor rather than plasma membrane fusion. In contrast, the HOP test has historically been the only functional assay to directly evaluate the ability of human sperm to interact and fuse with an oocyte membrane (Yanagimachi et al., 1976). In fact, parameters such as the hyaluronan binding assay (HBA) were shown not to correlate and to be non-redundant (Lazarevic et al., 2010). Even though the HOP test showed promising capacity to predict the outcomes of conventional IVF and ICSI (Freeman et al., 2001), over time it became obsolete (World Health Organization, HRP, 2021) mainly due to its elevated complexity, cost, and difficulty to standardize between laboratories. In this sense, hSPICER rises as an attractive alternative, as established cell lines can be maintained and distributed among diagnostic laboratories.

Altogether, this study presents an optimized cell-based methodology (overviewed in Figure 7) to directly assess human sperm fertilization competence with important implications for the quantitative diagnosis and potential treatments of male infertility.

**Figure 7:**
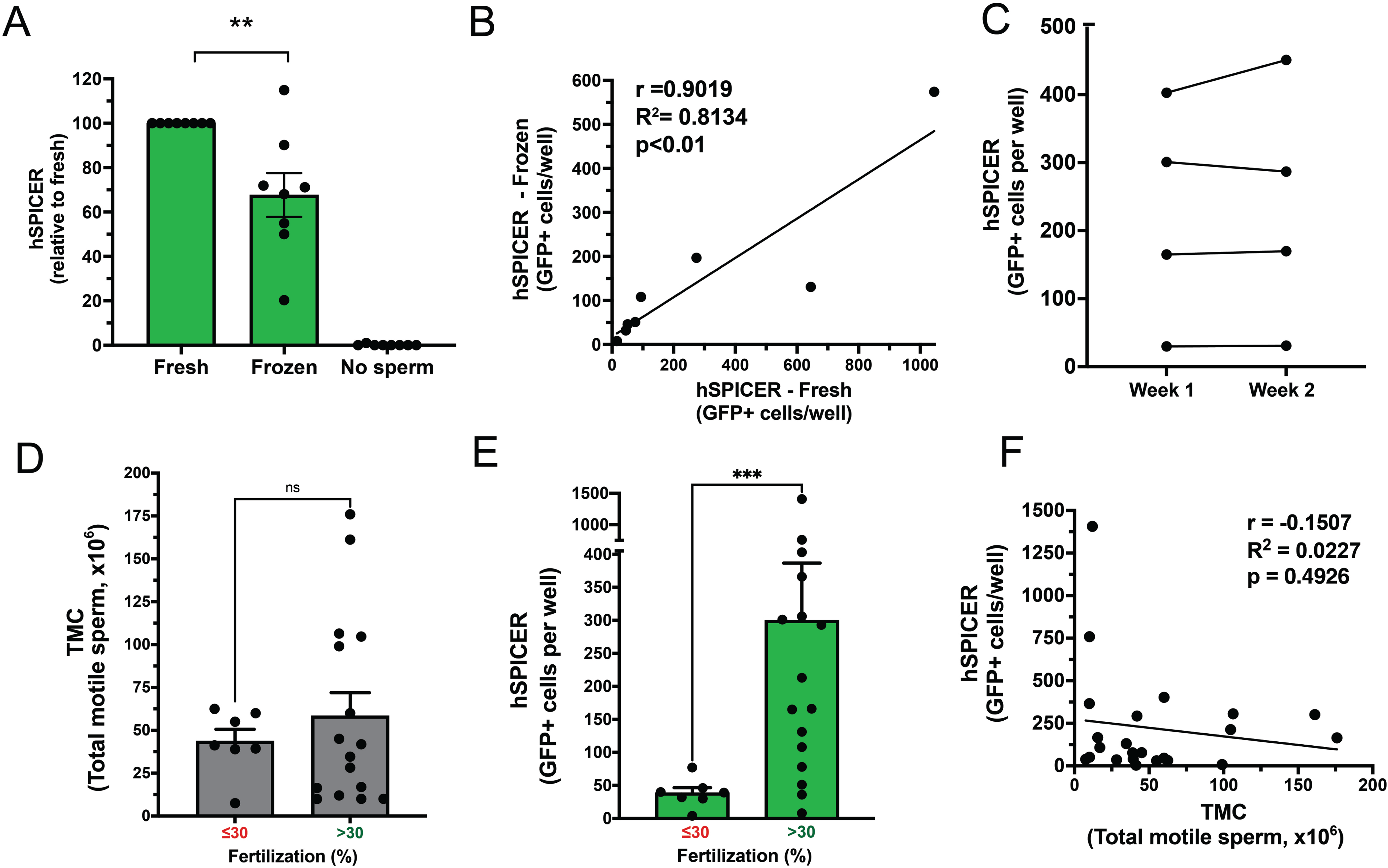
Overview of the optimized hSPICER workflow. (A) Generation of stable hSPICER reporter cell lines: HEK293T cells stably expressing DSP1-7 or DSP8-11 were generated by transfection and puromycin selection, followed by lentiviral transduction with hJUNO-flag::mCherry. (B) Sperm preparation and basic semen analysis: Following liquefaction, semen parameters were assessed according to WHO guidelines. Motile sperm were isolated by density-gradient centrifugation, washed, cryopreserved, and subsequently thawed and adjusted to the concentration required for the assay. (C) hSPICER assay: Reporter cell populations were mixed at a 1:1 ratio and seeded in poly-L-lysine-coated plates. After 24 h, sperm were added in fertilization medium and co-incubated with the reporter cells for 18 h. Cells were then washed, fixed, and sperm-induced cell–cell fusion was quantified as the number of GFP-positive cells per well.

## Materials and Methods

### Cell culture

BHK cells (Cat# CCL-10; ATCC, RRID: CVCL_1915) and HEK293T cells (Cat# CRL-3216; ATCC, RRID: CVCL_0063) were grown and maintained in DMEM containing 10% FBS and cultured at 37°C in 5% CO_2_. In all cases, the cell lines tested negative for mycoplasma contamination.

### Generation of tagged human JUNO constructs

In all cases, oligonucleotides from Sigma-Aldrich and FastDigest restriction enzymes (Thermo Fisher Scientific) were used, and all constructs were verified by DNA sequencing (Macrogen). For the tagging of human JUNO: (i) N-terminal flag-hJUNO was amplified using a forward primer containing the signal peptide sequence followed by the flag-tag sequence; (ii) hJUNO-C-terminal flag was amplified using a reverse primer containing the flag-tag sequence followed by the GPI-anchor signal sequence; and (iii) hJUNO-TMD-V5 was generated by overlap PCR, combining the ectodomain of JUNO with the transmembrane domain and V5 sequence from a human IZUMO1 plasmid available in the lab. All variants were digested with NheI and EcoRV and ligated into pCI::H2B-RFP previously digested with NheI and SmaI.

For lentiviral transduction, the hJUNO-C-terminal flag-sequence was cloned into pCSCW::IRES::mCherry by Gibson cloning. The complete list of primers and plasmids used in this study is shown in Table S1.

### Cell transfection

Freshly prepared plasmids were transiently transfected into cells using 2 μl jetPRIME (PolyPlus-transfection) per µg of DNA in 100 μl of reaction buffer for every ml of medium, unless otherwise specified. HEK293T cells stable lines for Dual Split Proteins (DSP) 1–7 and 8–11 were prepared by transfecting pIRESpuro3-DSP1–7 and pIRESpuro3-DSP8-11, respectively, and selecting with 2 µg/ml of puromycin for 10–13 days as previously described (Brukman et al., 2024; H. Wang et al., 2014). To establish HEK293T cells with stable expression of hJUNO, lentivirus was generated as previously described (Hindi et al., 2023). Briefly, 2.5×10^6^ HEK293T cells in 35 mm plates and 3 ml of medium were transfected with 10 μg pCSCW::hJUNO-flag::mCherry transfer plasmid, 7.5 μg psPAX2 packaging plasmid (Addgene, plasmid #12260), and 2.5 μg pMD2.G envelope plasmid (Addgene, plasmid #12259). 48 h later, the supernatant containing viral particles was collected, centrifuged at 2500 rpm for 5 min, and passed through a 0.45 μm filter. The viral suspension was diluted 1:1 with fresh medium, 6 μg/ml polybrene (Sigma) was added, and the transduction of HEK293T(DSP1-7) and HEK293T(DSP8-11) was carried out by incubation with the viral suspension overnight.

### Sperm preparation and basic analysis

All procedures involving human semen samples were conducted according to the recommendations of the World Health Organization (World Health Organization, HRP, 2021) and approved by the Helsinki Committees of HaEmek Medical Center (Afula, Israel, 0066-24EMC) and Clalit Health Services (Israel, 0139-23-CMC). All donors were provided with written information about the study before giving informed consent. After liquefaction, a basic semen analysis was conducted (World Health Organization, HRP, 2021): semen volume was measured immediately after liquefaction using a calibrated serological pipette and recorded in millilitres (ml); sperm concentration was determined using Makler counting chamber under phase-contrast microscopy at 200X magnification and expressed as millions per millilitre (10^6^/ml); morphological assessment was performed according to WHO criteria. The Total Motile Cells were calculated as the product of Volume (ml) x Sperm Concentration (million cells/ml) x Motility (%)/100. The semen parameters of all the normozoospermic donors fell within the WHO normality criteria. The sperm cells from both donors and patients were isolated by a 40%/80% gradient fraction of PureCeption® (Catalog #ART-2100, CooperSurgical, Inc., Trumbull, CT, USA) prepared in medium and centrifuged at 400 x g for 20 min at room temperature. The recovered pellet was resuspended in 5 ml of sperm washing medium (Catalog #9983, FUJIFILM Irvine Scientific, Santa Ana, CA, USA) and centrifuged at 600 x g for 10 min to remove residual gradient medium. The final pellet was resuspended in culture medium and assessed for sperm concentration, motility and morphology prior to IVF procedures. Then, sperm were diluted to a final concentration of 1×10^7^ cells/ml in fertilization medium: Human Tubal Fluid (HTF) medium (Quinn et al., 1985) supplemented with 4 mg/ml of Bovine Serum Albumin (Catalog # A7906, Sigma) or in SAGE 1-Step^TM^ medium (Catalog #67020060, CooperSurgical). When indicated, the cells were incubated for 18 h at 37°C in an atmosphere with 5% vol/vol CO_2_ in air to allow sperm capacitation (Battistone et al., 2013).

### Sperm cryopreservation and thawing

When indicated, the processed sperm cells were cryopreserved by mixing in a 1:1 ratio with a commercial Freezing Medium (Catalog #90128, FUJIFILM Irvine Scientific) following the manufacturer’s instructions. The cryovials were incubated at 4 °C for 1 h, then held at −20 °C for at least 20 min before transfer to liquid nitrogen for storage. When needed, the samples were thawed at room temperature and washed once with Multipurpose Handling Medium (Catalog #90166, FUJIFILM Irvine Scientific), centrifuged at 600 x g for 10 min, and resuspended in the fertilization medium.

### Sperm-to-BHK cell fusion

BHK cells were grown on 24-well glass-bottom tissue-culture plates. 24 h after plating, cells were transfected with 0.25 µg pcDNA3.1-EGFP-MBD-nls plasmids and 0.5 µg of either pCI::H2B-RFP or pCI::hJUNO::H2B-RFP. 24 h after transfection, 10^7^ capacitated sperm in HTF medium were added to each well and co-incubated with the BHK cells for 6 h at 37°C and 5% CO2. Then, the mitochondria were stained by incubation with 0.5 mM MitoTracker Deep Red FM (Catalog #M22426, Invitrogen) in PBS for 30 min at 37°C. After one wash with PBS, the cells were fixed with 4% PFA in PBS and stained with 1 µg/ml DAPI. Micrographs were obtained using wide-field illumination using an ELYRA system S.1 microscope (Plan-Apochromat 20X NA 0.8; Zeiss). The fusion of sperm was confirmed by evaluating the transfer of EGFP-MBD-nls signal from the BHK cell to the sperm nuclei.

### Sperm-induced cell-cell fusion (SPICER) assay: optimization using normozoospermic donors

The fusion of somatic cells was evaluated by content mixing assay using the Dual Split Protein system, as previously described (Brukman et al., 2024). Briefly, HEK293T cells stably expressing DSP1-7 or DSP8-11 (10^5^ each) were mixed and seeded on 24-well glass-bottom tissue-culture plates (Cellvis, Cat. No. P24-1.5H-N) pre-treated with 20 μg/ml of Poly-L-lysine and transfected with 1 µg pCI::H2B-RFP or pCI::hJUNO::H2B-RFP. 18 h after transfection, 10^7^ sperm cells in fertilization medium were added to the HEK293T cells and co-incubated for 18 h, after which they were washed once with Dulbecco’s Phosphate Buffered Saline containing calcium and magnesium, fixed with 4% PFA (Bar Naor, Cat. No. BN15710) in PBS at room temperature for 20 minutes, and stained with 1 µg/ml DAPI. When indicated, different time points and sperm concentrations were employed. Micrographs were obtained as mentioned above, using wide-field illumination using an ELYRA system S.1 microscope (Plan-Apochromat 20X, NA 0.8; Zeiss). For GFP we used Ex 470/40, Beamsplitter 493, Em 525/50. For mCherry and RFP we used Ex 550/25, Beamsplitter 570, Em 605/70. For DAPI we used Ex 365, Beamsplitter 395, Em 445/50. The number of GFP-positive cells per well was determined.

### Sperm-induced cell-cell fusion (SPICER) assay: evaluation of patients

HEK293T cells stably expressing hJUNO and DSP1-7 or DSP8-11 (10^5^ each) were mixed and seeded on 24-well glass-bottom tissue-culture plates (Cellvis, Cat. No. P24-1.5H-N) pre-treated with 20 μg/ml of Poly-L-lysine. 18 h after seeding, 2×10^6^ sperm cells (fresh or thawed) in fertilization medium were added to the HEK293T cells and co-incubated for 18 h, after which they were washed once with Dulbecco’s Phosphate Buffered Saline containing calcium and magnesium, fixed with 4% PFA (Bar Naor, Cat. No. BN15710) in PBS at room temperature for 20 minutes, and stained with 1 µg/ml DAPI. Micrographs were obtained as mentioned above and the number of GFP-positive cells per well was determined.

### Immunostaining and analysis of localization

HEK293T cells (1.5×10^5^ cells/well) were seeded in 24-well glass-bottom tissue-culture plates coated with human fibronectin (5 µg/cm²; Biological Industries, Cat. No. 05-750-1F). Twenty-four hours after plating, cells were transfected with 0.5 µg of one of the following plasmids: pCI::hJUNO-flag::H2B-RFP, pCI::flag-hJUNO::H2B-RFP, or pCI::hJUNO-TMD-V5::H2B-RFP, as described above, and incubated for an additional 24 h.

For non-permeabilized conditions, cells were fixed with 4% PFA for 10 min at room temperature. For permeabilization, cells were fixed with 4% PFA for 20 min at room temperature and subsequently permeabilized with 0.1% Triton X-100 in PBS for 5 min. In all cases, the cells were washed with PBS and incubated with mouse anti-flag antibody (1:1,000; Sigma-Aldrich, Cat. No. F3165) or mouse anti-V5 antibody (1:500; Invitrogen, Cat. No. R960-25) for 1 h at room temperature, followed by Alexa Fluor 488 donkey anti-mouse secondary antibody (1:500; Invitrogen, Cat. No. A-21202) for 1 h at room temperature. Cells were washed three times with PBS between antibody incubation steps. Images were acquired by wide-field fluorescence microscopy using an ELYRA PS.1 system equipped with a Plan-Apochromat 63×/1.4 NA objective (Zeiss) and an iXon EMCCD camera (Andor), using ZEN microscopy software (version 7.0.4.0; RRID: SCR_013672; Zeiss).

Following immunofluorescence staining, the percentages of cells expressing hJUNO and cells displaying cell-surface localization of hJUNO were quantified. Approximately 400 transfected cells were analyzed for each condition. The percentage of stained cells was calculated as the number of flag/V5-positive cells divided by the total number of transfected cells and multiplied by 100.

### Immunoblot analysis

HEK293T wild-type, HEK293T(DSP1-7) and HEK293T(DSP8-11) cells with stable expression of hJUNO-flag (7×10^5^ cells of each) were seeded in 6-well plates. 24 h later, cells were washed with PBS (without Ca^2+^ and Mg^2+^), then treated for 10 min with 0.75 ml of 0.05% EDTA solution, and transferred to a 1.5 ml tube. After a centrifugation (5 min at 400 g), the pellet was mixed with 100 μl of reducing sample buffer (#S3401; Sigma-Aldrich) and incubated 10 min at 95°C. After pelleting by centrifugation, samples were loaded on a BOLT 4–12% Bis-Tris Plus gel and transferred to polyvinylidene fluoride (PVDF) membrane for 2 h, membranes blocked (5% skim milk, Sigma-Aldrich Cat. No. 70166), and incubated with primary antibody anti-flag antibody (1:1,000; Sigma-Aldrich, Cat. No. F3165) or anti-tubulin (1:500, Sigma-Aldrich, Cat. No. T9026) overnight at 4°C, and HRP-conjugated goat anti-mouse secondary antibody (1:15,000, Cat# 115-035-003; Jackson ImmunoResearch Labs, RRID: AB_10015289) for 1 h at room temperature. Chemiluminescent signals were detected using an ECL substrate (#34078; Thermo Fisher Scientific); see Figures 5E and S1).

### Statistics and data analysis

Results are shown as means ± SEM. For each experiment, at least four independent biological repetitions were performed. The significance of differences between the averages was analyzed as described in the figure legends (GraphPad Prism 9, RRID: SCR_002798). ROC curve analysis was performed to evaluate the ability of hSPICER to discriminate between samples associated with <30% and ≥30% IVF fertilization. The AUC, sensitivity, and specificity were also calculated using GraphPad Prism. Figures were prepared with Photoshop CS6 (Adobe, RRID: SCR_014199), Illustrator CS6 (Adobe, RRID: SCR_010279), BioRender (RRID:SCR_018361), and FIJI (ImageJ 1.53 c, RRID: SCR_002285). Proofreading, grammar, and spelling were assisted by ChatGPT4 (OpenAI).

## Data Availability

The data underlying this article are available in the article and in its online supplementary material.

## Authors’ contributions

B.P. and N.G.B. conceived and supervised the study. N.G.B. designed and conducted experiments, processed the data, and prepared the figures. R.A. performed immunostainings and western blot and assisted with the preparation of figures. M.K., R.L., S.B., and R.B.F. conducted clinical assessment, obtained Helsinki ethical approvals, and recruited donors and patients. N.G.B and B.P. wrote the manuscript, which was reviewed by M.K., R.L. and R.B.F and approved by all authors.

## Acknowledgements

We thank Yoram Dekel for support in clinical assessment, expert advice, and obtaining Helsinki ethical approvals. Kazuo Yamagata (Kindai University, Higashiosaka City, Osaka, Japan) for the EGFP-MBD-NLS plasmid; Zene Matsuda (University of Tokyo, Japan) for the DSP plasmids; Gavin Wright and Enrica Bianchi (University of York, York, UK) for the plasmids encoding for human and hamster JUNO and human IZUMO1. Yael Iosilevskii, Yoram Dekel, and Dan Cassel for critically reading the manuscript, and the Brukman and Podbilewicz lab members for many enlightening discussions. This work is dedicated to the memory of Shira Baram.

## Supplemental Table Legends

**Table S1.** Plasmids and primers used for this study. Plasmids used in this study and their corresponding sources are listed. For constructs generated in this study, the parental plasmid and primers used for cloning are indicated.

**Table S2.** Semen parameters and hSPICER results of normozoospermic donors used for assay optimization. Semen volume (ml), sperm concentration (x10 /ml), total motility (%), total motile cell count (TMC), progressive motility (%), viability (%), and normal morphology (%) are shown for each donor. TMC (x10) was calculated as semen volume x sperm concentration x total motility/100. hSPICER results are expressed as the number of GFP-positive cells per well under the indicated experimental conditions: “Vector + Sperm” corresponds to control cells lacking hJUNO expression co-incubated with sperm; “JUNO (w/o sperm) corresponds to hJUNO-expressing cells incubated in the absence of sperm; and “JUNO + Sperm” corresponds to hJUNO-expressing cells co-incubated with sperm. Donor samples were classified as normozoospermic based on conventional semen analysis parameters.

**Table S3:** Semen parameters, IVF outcomes, and hSPICER values of patients undergoing conventional IVF treatment. Semen volume, sperm concentration, total motility, and Total Motile Cells (TMC) are shown for each sample. TMC (x10) was calculated as semen volume (ml) x sperm concentration (x10 /ml) x total motility (%)/100. Fertilization rate was calculated as the number of fertilized oocytes divided by the total number of inseminated oocytes x 100. The fertilization results were defined as high fertilization with >30%, whereas rates ≤30% were classified as low fertilization. hSPICER activity is expressed as the number of GFP-positive cells per well and was evaluated using fresh sperm when available and/or frozen-thawed aliquots of the same semen sample. The no-sperm condition was included as a negative control. For the patient with repeated IVF cycles (indicated by the same patient number followed by different letters), each row represents an independent semen sample and IVF cycle from the same patient; the average values used for patient-level analyses are shown in the corresponding “average” row. “–”, not determined; “n.a.”, not applicable.

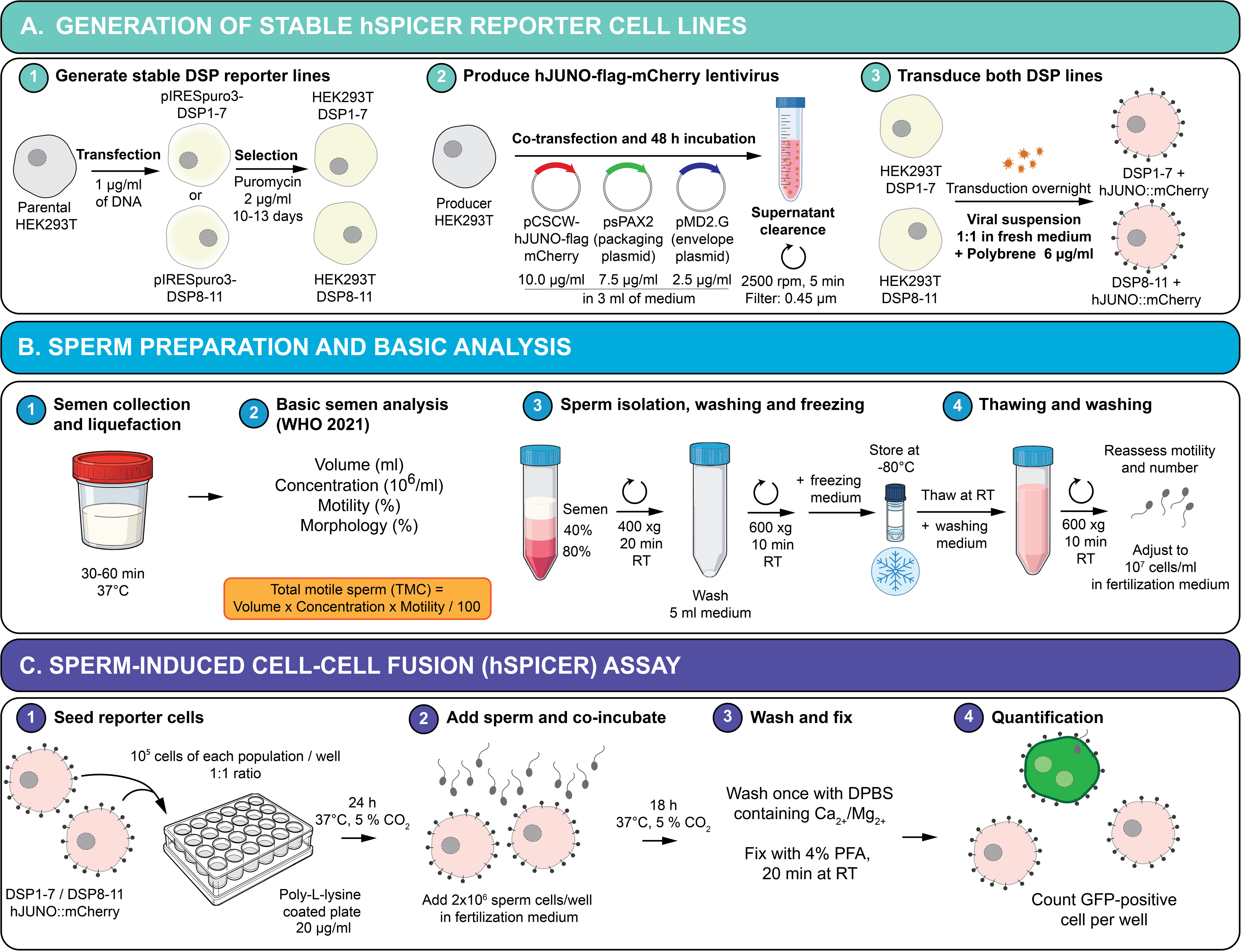

## References

1. Agarwal A, Baskaran S, Parekh N, Cho C-L, Henkel R, Vij S, Arafa M, Panner Selvam MK, Shah R. Male infertility. Lancet. 2021; 397:319–333.

2. Aitken RJ, Elton RA. Quantitative analysis of sperm-egg interaction in the zona-free hamster egg penetration test. Int J Androl. 1986; 9:14–30.

3. Allingham J, Floriano WB. Genetic diversity in the IZUMO1-JUNO protein-receptor pair involved in human reproduction. PLoS One. 2021; 16.

4. Avinoam O, Fridman K, Valansi C, Abutbul I, Zeev-Ben-Mordehai T, Maurer UE, Sapir A, Danino D, Grünewald K, White JM et al. Conserved eukaryotic fusogens can fuse viral envelopes to cells. Science. 2011; 332:589–592.

5. Barbaux S, Ialy-Radio C, Chalbi M, Dybal E, Homps-Legrand M, Do Cruzeiro M, Vaiman D, Wolf JP, Ziyyat A. Sperm SPACA6 protein is required for mammalian sperm-egg adhesion/fusion. Sci Rep. 2020; 10:5335.

6. Battistone MA, Da Ros VG, Salicioni AM, Navarrete FA, Krapf D, Visconti PE, Cuasnicú PS. Functional human sperm capacitation requires both bicarbonate-dependent PKA activation and down-regulation of Ser/Thr phosphatases by Src family kinases. Mol Hum Reprod. 2013; 19:570–580.

7. Bianchi E, Doe B, Goulding D, Wright GJ. Juno is the egg Izumo receptor and is essential for mammalian fertilization. Nature. 2014; 508:483–487.

8. Bianchi E, Jiménez-Movilla M, Cots-Rodríguez P, Viola C, Wright GJ. No evidence for a direct extracellular interaction between human Fc receptor-like 3 (MAIA) and the sperm ligand IZUMO1. Sci Adv. 2024; 10.

9. Boeri L, Belladelli F, Capogrosso P, Cazzaniga W, Candela L, Pozzi E, Valsecchi L, Papaleo E, Viganò P, Abbate C et al. Normal sperm parameters per se do not reliably account for fertility: a case-control study in the real-life setting. Andrologia. 2021; 53.

10. Brukman NG, Nakajima KP, Valansi C, Flyak K, Li X, Higashiyama T, Podbilewicz B. A novel function for the sperm adhesion protein IZUMO1 in cell-cell fusion. J Cell Biol. 2023; 222:e202207147.

11. Brukman NG, Valansi C, Podbilewicz B. Sperm induction of somatic cell-cell fusion as a novel functional test. Elife. 2024; 13:e94228.

12. Cots-Rodríguez P, Wang X, Sanchez-Tudela M, Siu KK, Yip P, Gómez E, Lee JE, Hamze JG, Jiménez-Movilla M. JUNO-coated beads as a functional assay to capture and characterize fertilization-competent human sperm. Hum Reprod Open. 2026; (1):hoag010.

13. de la Fuente D, Maroto M, Cajas YN, Canon-Beltran K, Fernandez-Gonzalez R, Munoz-Maceda A, Sanchez-Puig JM, Blasco R, Cots-Rodríguez P, Avilés M et al. Oviductin sets the species-specificity of the mammalian zona pellucida. Elife. 2025; 13.

14. Deneke VE, Blaha A, Lu Y, Suwita JP, Draper JM, Phan CS, Panser K, Schleiffer A, Jacob L, Humer T et al. A conserved fertilization complex bridges sperm and egg in vertebrates. Cell. 2024; 187:7066–7078.e22.

15. Duke LC, Cone AS, Sun L, Dittmer DP, Meckes DG Jr, Tomko RJ Jr. Tetraspanin CD9 alters cellular trafficking and endocytosis of tetraspanin CD63, affecting CD63 packaging into small extracellular vesicles. J Biol Chem. 2025; 301:108255.

16. Elofsson A, Han L, Bianchi E, Wright GJ, Jovine L. Deep learning insights into the architecture of the mammalian egg-sperm fusion synapse. Elife. 2024; 13.

17. Fernández JL, Cajigal D, López-Fernández C, Gosálvez J. Assessing sperm DNA fragmentation with the sperm chromatin dispersion test. Methods Mol Biol. 2011; 682:291–301.

18. Fernández JL, Muriel L, Goyanes V, Segrelles E, Gosálvez J, Enciso M, LaFromboise M, De Jonge C. Simple determination of human sperm DNA fragmentation with an improved sperm chromatin dispersion test. Fertil Steril. 2005; 84:833–842.

19. Fernández JL, Muriel L, Rivero MT, Goyanes V, Vazquez R, Alvarez JG. The sperm chromatin dispersion test: a simple method for the determination of sperm DNA fragmentation. J Androl. 2003; 24:59–66.

20. Freeman MR, Archibong AE, Mrotek JJ, Whitworth CM, Weitzman GA, Hill GA. Male partner screening before in vitro fertilization: preselecting patients who require intracytoplasmic sperm injection with the sperm penetration assay. Fertil Steril. 2001; 76:1113–1118.

21. Fujihara Y, Lu Y, Noda T, Oji A, Larasati T, Kojima-Kita K, Yu Z, Matzuk RM, Matzuk MM, Ikawa M. Spermatozoa lacking fertilization influencing membrane protein (FIMP) fail to fuse with oocytes in mice. Proc Natl Acad Sci U S A. 2020; 117:9393–9400.

22. Gervasi MG, Visconti PE. Chang’s meaning of capacitation: a molecular perspective. Mol Reprod Dev. 2016; 83:860–874.

23. Gierke T, Liu P, Lu Y, Tang S. Membrane fusions during mammalian fertilization. Adv Exp Med Biol. 2026; 1509:87–106.

24. González SN, Rebagliati Cid A, Sulzyk V, Weigel-Muñoz M, Santi CM, Visconti PE, Cuasnicu PS. CatSper-mediated sperm motility is required for mammalian gamete fusion. Sci Rep 2026, 16:29451.

25. Hamze JG, Sánchez JM, O’Callaghan E, McDonald M, Bermejo-Álvarez P, Romar R, Lonergan P, Jiménez-Movilla M. JUNO protein coated beads: a potential tool to predict bovine sperm fertilizing ability. Theriogenology. 2020; 155:168–175.

26. Hindi SM, Petrany MJ, Greenfeld E, Focke LC, Cramer AAW, Whitt MA, Khairallah RJ, Ward CW, Chamberlain JS, Prasad V et al. Enveloped viruses pseudotyped with mammalian myogenic cell fusogens target skeletal muscle for gene delivery. Cell. 2023; 186:2062–2077.e17.

27. Hirohashi N, Yanagimachi R. Sperm acrosome reaction: its site and role in fertilization. Biol Reprod. 2018; 99:127–133.

28. Holt WV. Basic aspects of frozen storage of semen. Anim Reprod Sci. 2000; 62:3–22.

29. Huszar G, Ozenci CC, Cayli S, Zavaczki Z, Hansch E, Vigue L. Hyaluronic acid binding by human sperm indicates cellular maturity, viability, and unreacted acrosomal status. Fertil Steril. 2003; 79 Suppl 3:1616–1624.

30. Inoue N, Hagihara Y, Wada I. Evolutionarily conserved sperm factors, DCST1 and DCST2, are required for gamete fusion. Elife. 2021; 10.

31. Inoue N, Ikawa M, Isotani A, Okabe M. The immunoglobulin superfamily protein Izumo is required for sperm to fuse with eggs. Nature. 2005; 434:234–238.

32. Ishikawa H, Meng F, Kondo N, Iwamoto A, Matsuda Z. Generation of a dual-functional split-reporter protein for monitoring membrane fusion using self-associating split GFP. Protein Eng Des Sel. 2012; 25:813–820.

33. Kaji K, Oda S, Shikano T, Ohnuki T, Uematsu Y, Sakagami J, Tada N, Miyazaki S, Kudo A. The gamete fusion process is defective in eggs of Cd9-deficient mice. Nat Genet. 2000; 24:279–282.

34. Kumar S, Valansi C, Haile MT, Li X, Flyak K, Dwivedy A, Abatiyow BA, Leeb AS, Kennedy SY, Camargo NM et al. Malaria parasites utilize two essential plasma membrane fusogens for gamete fertilization. Cell Mol Life Sci. 2022; 79:549.

35. Lamas-Toranzo I, Hamze JG, Bianchi E, Fernández-Fuertes B, Pérez-Cerezales S, Laguna-Barraza R, Fernández-González R, Lonergan P, Gutiérrez-Adán A, Wright GJ et al. TMEM95 is a sperm membrane protein essential for mammalian fertilization. Elife. 2020; 9.

36. Lazarevic J, Wikarczuk M, Somkuti SG, Barmat LI, Schinfeld JS, Smith SE. Hyaluronan binding assay (HBA) vs. sperm penetration assay (SPA): can HBA replace the SPA test in male partner screening before in vitro fertilization? J Exp Clin Assist Reprod. 2010; 7:2.

37. Le Naour F, Rubinstein E, Jasmin C, Prenant M, Boucheix C. Severely reduced female fertility in CD9-deficient mice. Science. 2000; 287:319–321.

38. Matsumura T, Noda T, Satouh Y, Morohoshi A, Yuri S, Ogawa M, Lu Y, Isotani A, Ikawa M. Sperm IZUMO1 is required for binding preceding fusion with oolemma in mice and rats. Front Cell Dev Biol. 2022; 9:810118.

39. Mio Y, Iwata K, Yumoto K, Kai Y, Sargant HC, Mizoguchi C, Ueda M, Tsuchie Y, Imajo A, Iba Y et al. Possible mechanism of polyspermy block in human oocytes observed by time-lapse cinematography. J Assist Reprod Genet. 2012; 29:951–956.

40. Miyado K, Yamada G, Yamada S, Hasuwa H, Nakamura Y, Ryu F, Suzuki K, Kosai K, Inoue K, Ogura A et al. Requirement of CD9 on the egg plasma membrane for fertilization. Science. 2000; 287:321–324.

41. Moi D, Nishio S, Li X, Valansi C, Langleib M, Brukman NG, Flyak K, Dessimoz C, de Sanctis D, Tunyasuvunakool K et al. Discovery of archaeal fusexins homologous to eukaryotic HAP2/GCS1 gamete fusion proteins. Nat Commun. 2022; 13:3880.

42. Molina LCP, Gunderson S, Riley J, Lybaert P, Borrego-Alvarez A, Jungheim ES, Santi CM. Membrane potential determined by flow cytometry predicts fertilizing ability of human sperm. Front Cell Dev Biol. 2020; 7:387.

43. Montag M, Köster M, van der Ven K, Bohlen U, van der Ven H. The benefit of artificial oocyte activation is dependent on the fertilization rate in a previous treatment cycle. Reprod Biomed Online. 2012; 24:521–526.

44. Murugesu S, Saso S, Jones BP, Bracewell-Milnes T, Athanasiou T, Mania A, Serhal P, Ben-Nagi J. Does the use of calcium ionophore during artificial oocyte activation demonstrate an effect on pregnancy rate? A meta-analysis. Fertil Steril. 2017; 108:468–482.e3.

45. Nakane S, Matsuda Z. Dual split protein (DSP) assay to monitor cell-cell membrane fusion. Methods Mol Biol. 2015; 1313:229–236.

46. Noda T, Blaha A, Fujihara Y, Gert KR, Emori C, Deneke VE, Oura S, Panser K, Lu Y, Berent S et al. Sperm membrane proteins DCST1 and DCST2 are required for sperm-egg interaction in mice and fish. Commun Biol. 2022; 5:332.

47. Noda T, Lu Y, Fujihara Y, Oura S, Koyano T, Kobayashi S, Matzuk MM, Ikawa M. Sperm proteins SOF1, TMEM95, and SPACA6 are required for sperm-oocyte fusion in mice. Proc Natl Acad Sci U S A. 2020; 117:11493–11502.

48. Oehninger S, Franken DR, Ombelet W. Sperm functional tests. Fertil Steril. 2014; 102:1528–1533.

49. Parrella A, Medrano L, Aizpurua J, Gómez-Torres MJ. Phospholipase C zeta in human spermatozoa: a systematic review on current development and clinical application. Int J Mol Sci. 2024; 25:1344.

50. Perez-Vargas J, Krey T, Valansi C, Avinoam O, Haouz A, Jamin M, Raveh-Barak H, Podbilewicz B, Rey FA. Structural basis of eukaryotic cell-cell fusion. Cell. 2014; 157:407–419.

51. Podbilewicz B, Leikina E, Sapir A, Valansi C, Suissa M, Shemer G, Chernomordik LV. The *C. elegans* developmental fusogen EFF-1 mediates homotypic fusion in heterologous cells and in vivo. Dev. Cell. 2006; 11:471–481.

52. Pregl Breznik B, Kovačič B, Vlaisavljević V. Are sperm DNA fragmentation, hyperactivation, and hyaluronan-binding ability predictive for fertilization and embryo development in in vitro fertilization and intracytoplasmic sperm injection? Fertil Steril. 2013; 99:1233–1241.

53. Quinn P, Kerin JF, Warnes GM. Improved pregnancy rate in human in vitro fertilization with the use of a medium based on the composition of human tubal fluid. Fertil Steril. 1985; 44:493–498.

54. Ruan JL, Liang SS, Pan JP, Chen ZQ, Teng XM. Artificial oocyte activation with Ca2+ ionophore improves reproductive outcomes in patients with fertilization failure and poor embryo development in previous ICSI cycles. Front Endocrinol (Lausanne). 2023; 14:1244507.

55. Sanyal D, Arya D, Nishi K, Balasinor N, Singh D. Clinical utility of sperm function tests in predicting male fertility: a systematic review. Reprod Sci. 2024; 31:863–882.

56. Satouh Y, Inoue N, Ikawa M, Okabe M. Visualization of the moment of mouse sperm-egg fusion and dynamic localization of IZUMO1. J Cell Sci. 2012; 125:4985–4990.

57. Valansi C, Moi D, Leikina E, Matveev E, Graña M, Chernomordik LV, Romero H, Aguilar PS, Podbilewicz B. Arabidopsis HAP2/GCS1 is a gamete fusion protein homologous to somatic and viral fusogens. J Cell Biol. 2017; 216:571–581.

58. Wang H, Li X, Nakane S, Liu S, Ishikawa H, Iwamoto A, Matsuda Z. Co-expression of foreign proteins tethered to HIV-1 envelope glycoprotein on the cell surface by introducing an intervening second membrane-spanning domain. PLoS One. 2014; 9.

59. Wang Y, Li R, Yang R, Zheng D, Zeng L, Lian Y, Zhu Y, Zhao J, Liang X, Li W et al. Intracytoplasmic sperm injection versus conventional in-vitro fertilisation for couples with infertility with non-severe male factor: a multicentre, open-label, randomised controlled trial. Lancet. 2024; 403:924–934.

60. World Health Organization. Infertility prevalence estimates, 1990-2021. 2023. World Health Organization, Geneva, Switzerland.

61. World Health Organization, Human Reproduction Programme. WHO laboratory manual for the examination and processing of human semen. 6th edn, 2021. World Health Organization, Geneva, Switzerland.

62. Yamagata K, Yamazaki T, Yamashita M, Hara Y, Ogonuki N, Ogura A. Noninvasive visualization of molecular events in the mammalian zygote. Genesis. 2005; 43:71–79.

63. Yanagimachi R. Mammalian fertilization. In: Knobil E, Neill JD, eds. The Physiology of Reproduction. 2nd edn, 1994. Raven Press, New York, pp. 189–317.

64. Yanagimachi R, Yanagimachi H, Rogers BJ. The use of zona-free animal ova as a test-system for the assessment of the fertilizing capacity of human spermatozoa. Biol Reprod. 1976; 15:471–476.

65. Young S, Schiffer C, Wagner A, Patz J, Potapenko A, Herrmann L, Nordhoff V, Pock T, Krallmann C, Stallmeyer B et al. Human fertilization in vivo and in vitro requires the CatSper channel to initiate sperm hyperactivation. J Clin Invest. 2024; 134.

