## Supplementary material for "Human Sperm-Induced Cell-Cell Fusion Requiring JUNO (hSPICER): A paradigm shift to test sperm fertilizing potential": Table S1

**Table S1. Plasmids and primers used for this study**

Plasmids used in this study and their corresponding sources are listed. For constructs generated in this study, the parental plasmid and primers used for cloning are indicated.

| Plasmid | Origin / Primers for cloning (5’-3’) |
|---|---|
| pCI::H2B-RFP | Williams et al., 2018, Addgene #92398 |
| pIRESpuro3-DSP1–7 | Wang et al., 2014 |
| pIRESpuro3-DSP8–11 | Wang et al., 2014 |
| pcDNA3.1-EGFP-MBD-nls | Yamagata et al., 2005 |
| pCI::human JUNO::H2B-RFP | Brukman et al., 2023 |
| pCI::mouse JUNO::H2B-RFP | Brukman et al., 2024 |
| pCI::hamster JUNO::H2B-RFP | Brukman et al., 2024 |
| pCMV6-human IZUMO1 | Origene #RC206671 |
| pCI::human IZUMO1-V5::H2B-RFP | This paper, from pCMV6-human IZUMO1.<br>Fwd primer (NheI): TTATCGCTAGCATGGGGCCGCATTTTACCCTC<br>Rvs primer (EcoRV and V5):<br>CCAGGATATCTCACGTAGAATCGAGACCGAGGAGAGGGTTAGGGATAGGCTTACCACGCG TTTGTTGCCTCGAATCTG |
| pCI::hJUNO-flag::H2B-RFP | This paper, from pCI::human JUNO::H2B-RFP.<br>Fwd primer (NheI): TTATCGCTAGCATGGCATGCTGGTGGCCGCTC<br>Rvs priemer (EcoRV + flag + GPI signal):<br>TCGATATCAGGAAAGGAACGGCAGGAACAGGGAGCAGACCATGATGGTGTAGGACAGTTC CCAGGATGGGGCAGAGCTCTTGTCATCGTCGTCCTTGTAGTCGGCGAAGAGGCGGGCCA C |
| pCI::flag-hJUNO::H2B-RFP | This paper, from pCI::human JUNO::H2B-RFP.<br>Fwd primer (NheI + signal peptide + flag):<br>TAAACGCTAGCATGGCATGCTGGTGGCCGCTCCTGCTAGAGCTGTGGACAGTCATGCCCA CCTGGGCTGGGGACTACAAGGACGACGATGACAAGGACGAGCTGCTCAACATCTG<br>Rvs primer (EcoRV): CCAGGATATCTCAGGAAAGGAACGGCAGGAAC |
| pCI::hJUNOTMD-V5::H2B-RFP | This paper, from pCI::human JUNO::H2B-RFP and pCI::human IZUMO1-V5::H2B-RFP |

| | |
|---|---|
| | Fwd primer (NheI): TTATCGCTAGCATGGCATGCTGGTGGCCGCTC<br>Inner Rvs primer:<br>GCGGCTTGCCAGCATTTTCTCGCTGGCGAAGAGGCGGGCCAC<br>Inner Fwd primer: GAGAAAATGCTGGCAAGCCGC<br>Rvs primer (EcoRV):<br>CCAGGATATCTCACGTAGAATCGAGACCGAGGAGAGGGTTAGGGATAGGCTTACCACGCG<br>TTTGTTGCCTCGAATCTG |
| pCSCW::IRES::mCherry | CSCW-mCherry, MGH Vector Core Facility |
| pCSCW::hJUNO-flag::mCherry | This paper, from pC::hJUNO-flag::H2B-RFP.<br>Fwd Gibson primer:<br>GTTTAGTGAACCGTCAGATCCGCTAGTCGAGCTCAAGCTTATGGCATGCTGGTGGCCGCT<br>C<br>Rvs Gibson primer:<br>GTAACGTTAGGGGGGGGGGAGGGAGAGGGGCGGATCAATTTCAGGAAAGGAACGGCAG<br>GAAC |
| psPAX2 | Addgene #12260 |
| pMD2.G | Addgene #12259 |
