## Supplementary material for "Human Sperm-Induced Cell-Cell Fusion Requiring JUNO (hSPICER): A paradigm shift to test sperm fertilizing potential": Table S2

**Table S2. Semen parameters and hSPICER results of normozoospermic donors used for assay optimization**

Semen volume (ml), sperm concentration (x$10^6$/ml), total motility (%), total motile cell count (TMC), progressive motility (%), viability (%), and normal morphology (%) are shown for each donor. TMC (x$10^6$) was calculated as semen volume x sperm concentration x total motility/100. hSPICER results are expressed as the number of GFP-positive cells per well under the indicated experimental conditions: “Vector + Sperm” corresponds to control cells lacking hJUNO expression co-incubated with sperm; “JUNO (w/o sperm) corresponds to hJUNO-expressing cells incubated in the absence of sperm; and “JUNO + Sperm” corresponds to hJUNO-expressing cells co-incubated with sperm. Donor samples were classified as normozoospermic based on conventional semen analysis parameters.

| Donor # | Date | Sperm analysis | | | | | | | SPICER (GFP+ cells per well) | | |
|---|---|---|---|---|---|---|---|---|---|---|---|
| | | Volume (mL) | Sperm concentration (10^6/mL) | Total Motility (%) | Total Motile Cells (TMC, 10^6) | Progressive motility (%) | Viability (%) | Normal morphology (%) | Vector + Sperm | JUNO (w/o sperm) | JUNO + Sperm |
| 1 | 20/12/2023 | 4.1 | 33 | 89 | 120.4 | 85 | 92 | 11 | 2 | 1 | 1853 |
| 2 | 03/01/2024 | 3.7 | 110 | 82 | 334.7 | 79 | 92 | 14 | 0 | 0 | 65 |
| 3 | 07/02/2024 | 3.2 | 30 | 53 | 50.0 | 49 | 83 | 10 | 1 | 1 | 314 |
| 4 | 21/03/2024 | 5.7 | 141 | 86 | 692.7 | 79 | 93 | 26 | 4 | 5 | 1585 |
| 5 | 03/04/2024 | 4.7 | 28 | 72 | 96.1 | 66 | 78 | 9 | 0 | 0 | 129 |
| 6 | 04/09/2024 | 1.4 | 21 | 91 | 26.8 | 85 | 97 | 18 | 0 | 0 | 132 |
| 7 | 09/09/2024 | 1.5 | 53 | 70 | 55.7 | 67 | 95 | 19 | 0 | 0 | 191 |
