## Supplementary material for "Human Sperm-Induced Cell-Cell Fusion Requiring JUNO (hSPICER): A paradigm shift to test sperm fertilizing potential": Table S3

**Table S3: Semen parameters, IVF outcomes, and hSPICER values of patients undergoing conventional IVF treatment**

Semen volume, sperm concentration, total motility, and Total Motile Cells (TMC) are shown for each sample. TMC ($x10^6$) was calculated as semen volume (ml) x sperm concentration ($x10^6$/ml) x total motility (%)/100. Fertilization rate was calculated as the number of fertilized oocytes divided by the total number of inseminated oocytes x 100. The fertilization results were defined as high fertilization with >30%, whereas rates ≤30% were classified as low fertilization. hSPICER activity is expressed as the number of GFP-positive cells per well and was evaluated using fresh sperm when available and/or frozen-thawed aliquots of the same semen sample. The no-sperm condition was included as a negative control. For the patient with repeated IVF cycles (indicated by the same patient number followed by different letters), each row represents an independent semen sample and IVF cycle from the same patient; the average values used for patient-level analyses are shown in the corresponding “average” row. “–”, not determined; “n.a.”, not applicable.

| Patient # | Date | Sperm analysis | | | | IVF results | | | | SPICER (GFP+ cells per well) | | | |
|---|---|---|---|---|---|---|---|---|---|---|---|---|---|
| | | Volume (mL) | Sperm concentration (10^6/ml) | Total Motility (%) | Total Motile Cells (TMC, 10^6) | Fertilized eggs | Total inseminated eggs | % of fertilization | High fertilization? (>30%) | Fresh sperm | Control (no sperm) | Frozen sperm (Week 1) | Frozen sperm (Week 2) |
| 1 | 15/12/2024 | 1 | 50 | 20 | 10.0 | 1 | 1 | 100.0 | Yes | 75 | 0 | 51 | - |
| 2 | 2/1/2025 | 0.5 | 30 | 50 | 7.5 | 0 | 3 | 0.0 | No | - | 0 | 39 | - |
| 3 | 2/1/2025 | 3 | 40 | 50 | 60.0 | 0 | 10 | 0.0 | No | 51 | 0 | 46 | - |
| 4 | 2/1/2025 | 3 | 100 | 50 | 150.0 | - | - | - | n.a. | 274 | 0 | 197 | - |
| 5 | 12/01/25 | 3.5 | 30 | 33 | 34.7 | 1 | 2 | 50.0 | Yes | 645 | 0 | 131 | - |
| 6 | 12/01/2025 | 2.5 | 50 | 50 | 62.5 | 1 | 7 | 14.3 | No | 45 | 0 | 32 | - |
| 7 | 14/01/2025 | 3.5 | 70.0 | 60 | 147.0 | - | - | - | n.a. | 1045 | 1 | 574 | - |
| 8 (average) | - | 1.5 | 33.3 | 33 | 15.6 | 19 | 23 | 82.6 | Yes | - | 0 | 166 | - |
| 8a | 21/01/2025 | 2 | 34 | 29 | 19.7 | 4 | 4 | 100.0 | Yes | - | 0 | 80 | - |

| | | | | | | | | | | | | | |
|---|---|---|---|---|---|---|---|---|---|---|---|---|---|
| 8b | 20/02/25 | 1.5 | 30 | 20 | 9.0 | 10 | 11 | 90.9 | Yes | - | 0 | 152 | - |
| 8c | 18/05/2025 | 1 | 36 | 50 | 18.0 | 5 | 8 | 62.5 | Yes | - | 0 | 266 | - |
| 9 | 21/01/25 | 0.5 | 51 | 39 | 9.9 | 6 | 15 | 40.0 | Yes | - | 0 | 366 | - |
| 10 | 2/2/25 | 3.5 | 26 | 31 | 28.2 | 8 | 9 | 88.9 | Yes | - | 0 | 36 | - |
| 11 | 18/02/25 | 2.5 | 90 | 44 | 99.0 | 1 | 2 | 50.0 | Yes | 16 | 0 | 8 | - |
| 13 | 20/02/25 | 5.5 | 15 | 50 | 41.3 | 10 | 49 | 20.4 | No | - | 0 | 4 | - |
| 14 | 6/3/2025 | 2 | 75 | 40 | 60.0 | 3 | 4 | 75.0 | Yes | - | 0 | 403 | 451 |
| 15 | 6/3/2025 | 2.5 | 50 | 44 | 55.0 | 1 | 8 | 12.5 | No | - | 0 | 30 | 31 |
| 16 | 18/03/2025 | 2 | 130 | 62 | 161.2 | 5 | 8 | 62.5 | Yes | - | 0 | 301 | 287 |
| 17 | 23/03/25 | 5 | 80 | 44 | 176.0 | 5 | 7 | 71.4 | Yes | - | 0 | 165 | 170 |
| 18 | 03/04/2025 | 3.5 | 65 | 46 | 104.7 | 5 | 7 | 71.4 | Yes | - | 0 | 213 | - |
| 19 | 06/04/2025 | 1.5 | 60 | 50 | 45.0 | 6 | 6 | 100.0 | Yes | - | 0 | 78 | - |
| 21 | 06/04/2025 | 0.6 | 50 | 40 | 12.0 | 2 | 2 | 100.0 | Yes | - | 0 | 1407 | - |

| | | | | | | | | | | | | | |
|---|---|---|---|---|---|---|---|---|---|---|---|---|---|
| 22 | 04/05/2025 | 7 | 40 | 38 | 106.4 | 3 | 9 | 33.3 | Yes | - | 0 | 306 | - |
| 23 | 13/05/2025 | 1 | 77 | 22 | 16.9 | 3 | 4 | 75.0 | Yes | 94 | 0 | 108 | - |
| 25 | 25/05/2025 | 1 | 50 | 20 | 10.0 | 3 | 4 | 75.0 | Yes | - | 0 | 759 | - |
| 26 | 29/05/2025 | 7 | 23 | 26 | 41.9 | 6 | 19 | 31.6 | Yes | - | 0 | 293 | - |
| 27 | 29/05/2025 | 2.5 | 45 | 35 | 39.4 | 0 | 1 | 0.0 | No | - | 0 | 40 | - |
| 28 | 29/05/2025 | 4 | 75 | 13 | 39.0 | 0 | 3 | 0.0 | No | - | 0 | 77 | - |
